# Light-dependent synthesis of a nucleotide second messenger controls bacterial motility

**DOI:** 10.1101/2021.07.06.451194

**Authors:** Jun Xu, Nobuo Koizumi, Yusuke V. Morimoto, Ryo Ozuru, Toshiyuki Masuzawa, Shuichi Nakamura

## Abstract

Nucleotide second messengers are universally crucial factors for the signal transduction of various organisms. In prokaryotes, cyclic nucleotide messengers are involved in the bacterial life cycle and function, such as virulence, biofilm formation, and others mainly via gene regulation. Here we show that the swimming motility of a soil bacterium is rapidly modulated by cyclic adenosine monophosphate (cAMP) synthesized upon light exposure. Analysis of a loss-of-photoresponsivity mutant obtained by transposon random mutagenesis determined the novel sensory gene, and its expression in *Escherichia coli* through codon optimization revealed the light-dependent synthesis of cAMP. GFP labeling showed the localization of the photoresponsive enzyme at the cell poles where flagellar motors reside. The present findings highlight the new role of cAMP that rapidly controls the flagella-dependent bacterial motility and the global distribution of the discovered photoactivated cyclase among diverse microbial species.

---

In all domains of life, nucleotide messengers cAMP, cGMP, c-di-AMP, and c-di-GMP have vital roles in signaling networks, allowing organisms to recognize changes in environmental factors and regulate their physiologies and behaviors. Many cyclic nucleotide-binding proteins have been identified, e.g., the binding of cAMP or cGMP to cyclic nucleotide-gated channels triggers cation flux^1^. The capacitation of mammalian sperms involves phosphorylation of the associated proteins by cAMP-dependent activation of protein kinase A, and its flagellar beating is accelerated by cAMP derived from soluble adenylyl cyclase^2^. Also, the c-di-GMP level is known to be involved in biofilm formation, motility, and virulence of bacteria^3,4^.

Here we show that the bacterial motility is rapidly modulated by cAMP generated upon light exposure. Motile bacteria possess varied machinery for moving in liquid or over surfaces^5^. A major motility form is flagella-dependent swimming: The peritrichous bacteria *Escherichia coli* and *Salmonella enterica* swim by rotating bundled flagella; *Vibrio cholerae* and *Pseudomonas aeruginosa* are propelled by a single polar flagellum^6^. While these species possess flagella at the cell exterior, spirochetes such as *Borrelia burgdorferi* (Lyme disease pathogen) and *Treponema pallidum* (syphilis) have their flagella (endoflagella) within the periplasmic space^7^. The endoflagella of spirochetes are thought to rotate within the periplasmic space, rolling or transforming the spiral cell body for generating thrust^7^. We discovered that the spirochete *Leptospira kobayashii* isolated from soil in Japan^8,9^ drastically alters the swimming pattern immediately after sensing light. We demonstrated the responsibility of cAMP synthesized by a novel photoresponsive adenylyl cyclase for motility control. It has been known that cAMP affects gene regulation for pili synthesis in cyanobacteria^10–12^, but the soil bacteria respond to light exposure in subsecond order. We termed the novel adenylyl cyclase as *L. kobayashii* photoactivation-associated adenylyl cyclase (LkPAAC). This is the first report of cAMP-dependent rapid control of flagella-driven bacterial motility.

## Results

### Light modulates swimming of soil bacteria

The genus *Leptospira* possesses two endoflagella (one flagellum per cell end, Supplementary Fig. 1). *Leptospira* spp. swim by rotating the coiled cell body (swimming mode), and smooth swimming is frequently intermitted by rotation without migration (rotation mode)^13,14^. It has been known that the transition between swimming and rotation modes of *Leptospira* spp. occurs stochastically, and that the switching frequency is affected by viscosity^15^ and chemical substrates^16,17^. In contrast, we found that *L. kobayashii* exclusively showed rotation in the dark (~ 10 lux, “Dark” in Fig. 1), but their smooth swimming is triggered by light exposure (~ 1300 lux, “Bright” in Fig. 1, Movie 1, Supplementary Fig. 2). Since light stimulation increased the migration distance of the bacteria by the directed swimming (Supplementary Fig. 3), the light-dependent motility was quantified using swimming velocity of individual cells (i.e., migration distance per second), showing an increase of velocity up to four-fold by light (Fig. 1b, Supplementary Fig. 2). The smooth swimming reached the maximum velocity at ~ 1 s after stimulation (Fig. 1c). The light-responsivity depends on the light intensity, and the duration time for unidirectional swimming got longer with the increase of light intensity (Supplementary Fig 2). The response to light was observed even at ~ 40 lux, which is less than 1/10 of a conventional experimental room illuminated by 32 W fluorescent lamps (500 – 600 lux) (Supplementary Fig 2). The bacteria can respond to green and blue lights but not to red (Fig 1d). Cumulative cell fractions obtained from the velocity histograms show that green light induces more cells to swim smoothly than blue light (Fig 1e).

**Figure 1.**
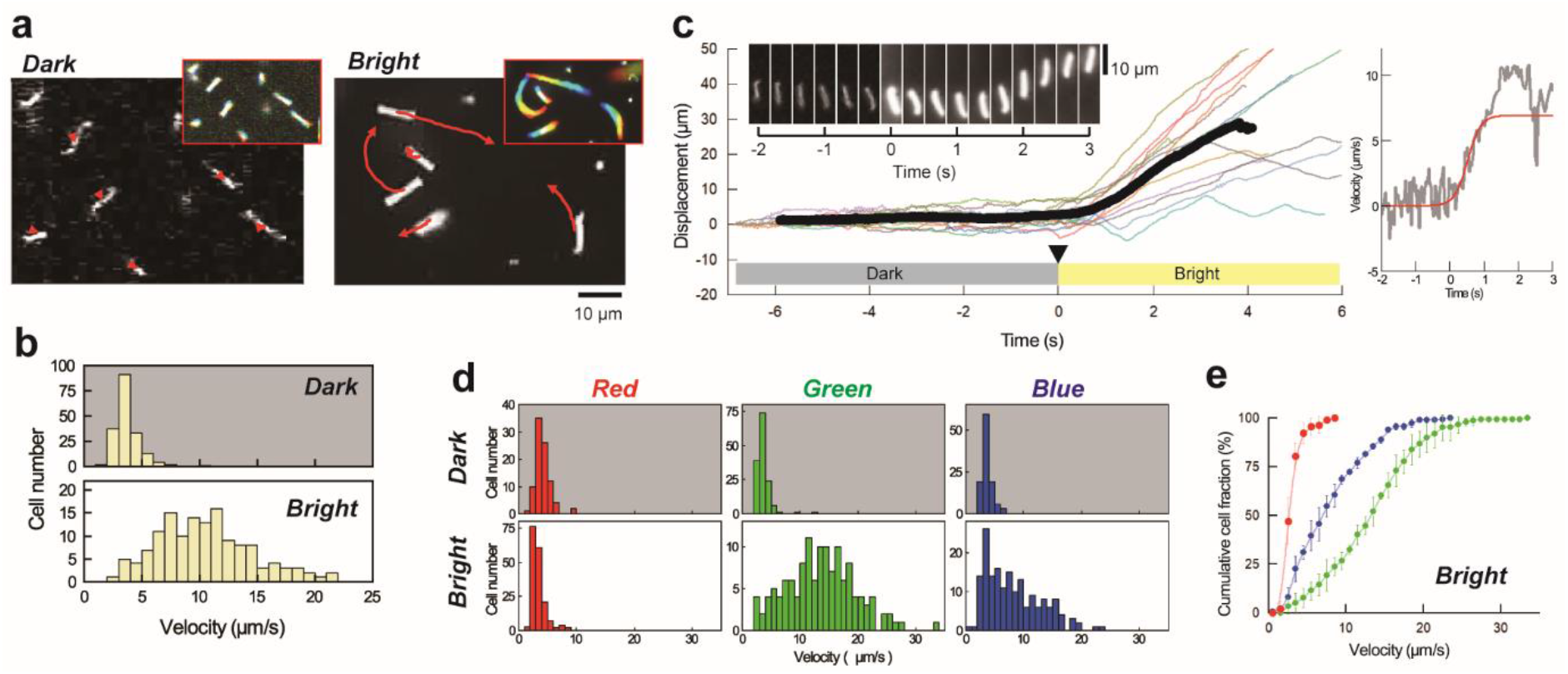
Light stimulation induces unidirectional swimming of spirochetal bacteria. **(a)** Swimming trajectories of *Leptospira kobayashii* cells under low-intensity (termed as “Dark” in this study) and high-intensity (termed as “Bright”) illumination. The results of 2-s tracking are shown. Colored footprints (inset, time courses in the order from red to blue) show almost zero net migration in dark. Note that the bacteria retain vigorous motility in dark (see the scene “Light OFF” in Movie1). As a light source, a halogen lamp was used with a wide band-pass filter (ca. 400-700 nm). **(b)** Velocities of individual cells measured in dark (n = 189 cells) and in bright (n = 150 cells). The data were obtained by three independent experiments. **(c)** Time courses of cell displacement. Example traces obtained from 14 cells (thin colored lines) and the averaged trace (thick black line) are shown. Neutral density filters were removed at the time point (indicated by the triangle and set as 0 s) so that the cells were suddenly exposed to stronger light. The sequential micrographs (inset) show the light-dependent initiation of directional swimming. The right panel shows the time course of velocity (gray) obtained from the mean displacement and the result of sigmoidal curve fitting (red). **(d)** Color dependence of the light-controlled motility. Band-pass filters with center wavelengths of 650 nm, 550 nm, and 488 nm were used for red, green, and blue illumination, respectively. The experiments were repeated three times, and ca. 100 cells were measured in total in each condition. **(e)** Cumulative cell fractions calculated from the histograms of bright shown in ***d***. The colors indicate those used for illumination: average values (closed circles) and standard deviations (error bars) of three independent experiments.

### Identification of the photosensor

To identify the sensor gene responsible for the *L. kobayashii* photoresponsivity, we explored loss-of-function mutants from a kanamycin-resistant library made by transposon random mutagenesis. One of ~ 2400 clones retained motility but lacked photoresponsivity (Fig 2a, Movie 2). The mutant carried a transposon insertion into the gene LPTSP3_g09850 (Fig. 2b, annotated as glycosyl hydrolase family), but the lost photoresponsivity was not restored to the mutant by the complementation of the gene (Supplementary Fig 4). Noting that the gene LPTSP3_g09840 encodes adenylate/guanylate cyclase domain-containing protein and is located immediately downstream of LPTSP3_g09850, we complemented 1H6 by both LPTSP3_g09850 and LPTSP3_g09840 genes, resulting in the recovery of photoresponsivity (Fig 2c, d). The photoresponsivity of 1H6 was recovered to the wild-type level by single complementation of LPTSP3_g09840 gene (Fig 2d), indicating that the gene is responsible for the photoresponsivity of *L. kobayashii*.

**Figure 2.**
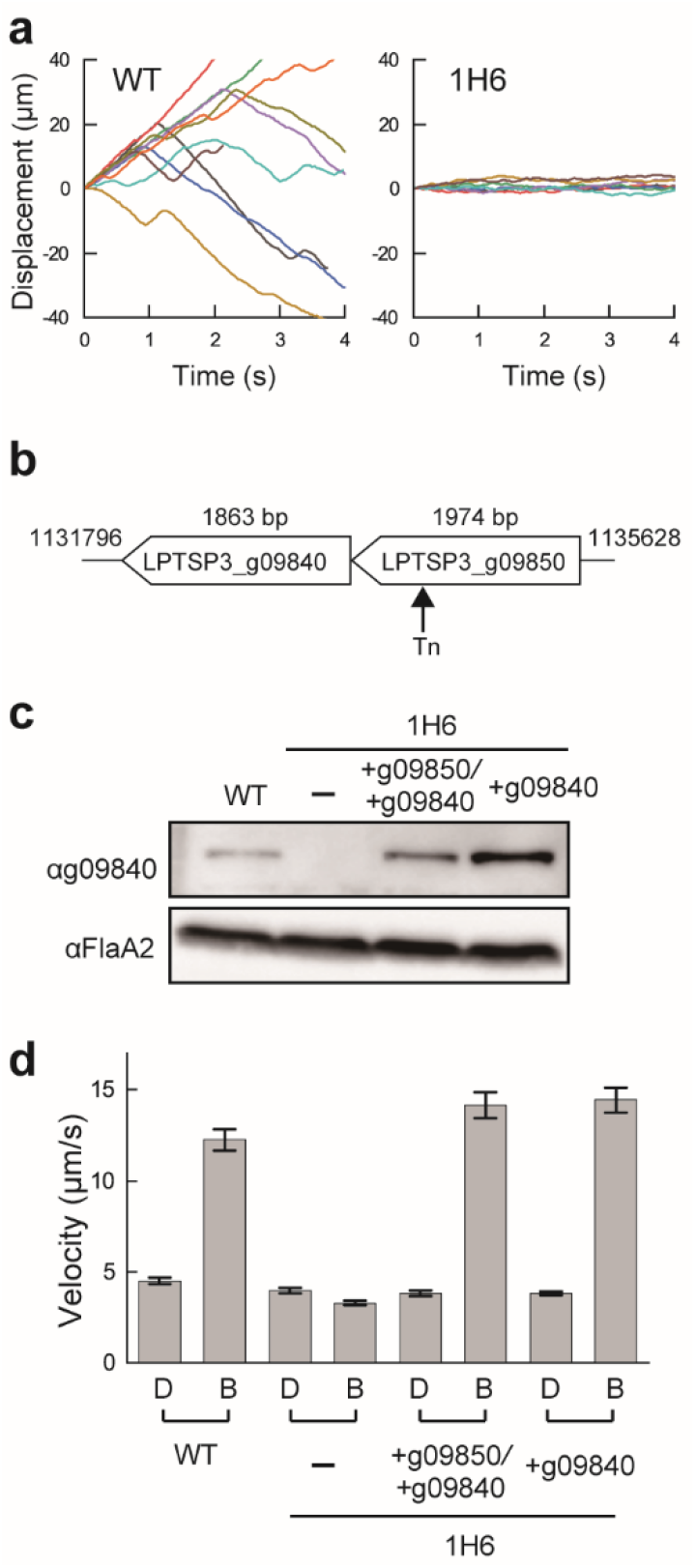
Identification of a photosensor in the spirochete *Leptospira*. (a) Time traces of the cell displacement under light exposure. The traces of 8 cells are shown in each strain. (b) Transposon insertion in the photoresponsivity-deficient mutant 1H6. The position of the transposon insertion is between 1133909th and 1133920th nucleotides of the *L. kobayashii* chromosome 1 (AP025028). (c) Immunoblotting of whole-cell lysates from the wild-type *L. kobayashii*, 1H6, and its complemented strains using anti-LkPAAC and anti-FlaA2 (control) antisera. (d) Effect of transposon insertion and gene complementation on photoresponsivity. The average values and standard errors were determined by three independent experiments (ca. 100 cells were measured in total in each condition).

### Light-dependent sensor activity

Photoactivated adenylyl cyclases (PACs) have been found in both eukaryotes and prokaryotes^18–24^. The alignment of BDA78054 (LPTSP3_g09840 gene product) with known functional PACs showed that the C-terminal region of BDA78054 contains a domain similar to the AC domain of known PACs, whereas the putative sensor domain of BDA78054 was distinct from the conventional PAC sensor BLUF or LOV domain (Supplementary Fig. 5). Since the light-dependent elevation of cAMP concentration was not detected in *L. kobayashii* cells (Supplementary Fig. 6a), we measured the enzyme activity using the overexpression system of *E. coli* carrying the codon-optimized LPTSP3_g09840 gene (Fig. 3a). The overexpressed BDA78054 showed the increase of the cAMP concentration in *E. coli* by light exposure (Fig. 3b). Very little cGMP was detected independently of light, which could be endogenous substrates (Supplementary Fig. 6b). Based on that the light-dependent cAMP synthesis, BDA78054 was named photoactivation-associated adenylyl cyclase (LkPAAC). The 1H6 mutant cells keep rotating in bright, but the addition of a membrane-permeable cAMP (8-Bromo-cAMP) induced smooth swimming of the mutant as well as the wild-type cells exposed to light (Fig. 3c). These results suggest that the observed modulation of flagellar rotation was induced by cAMP synthesized upon light stimulation.

**Figure 3.**
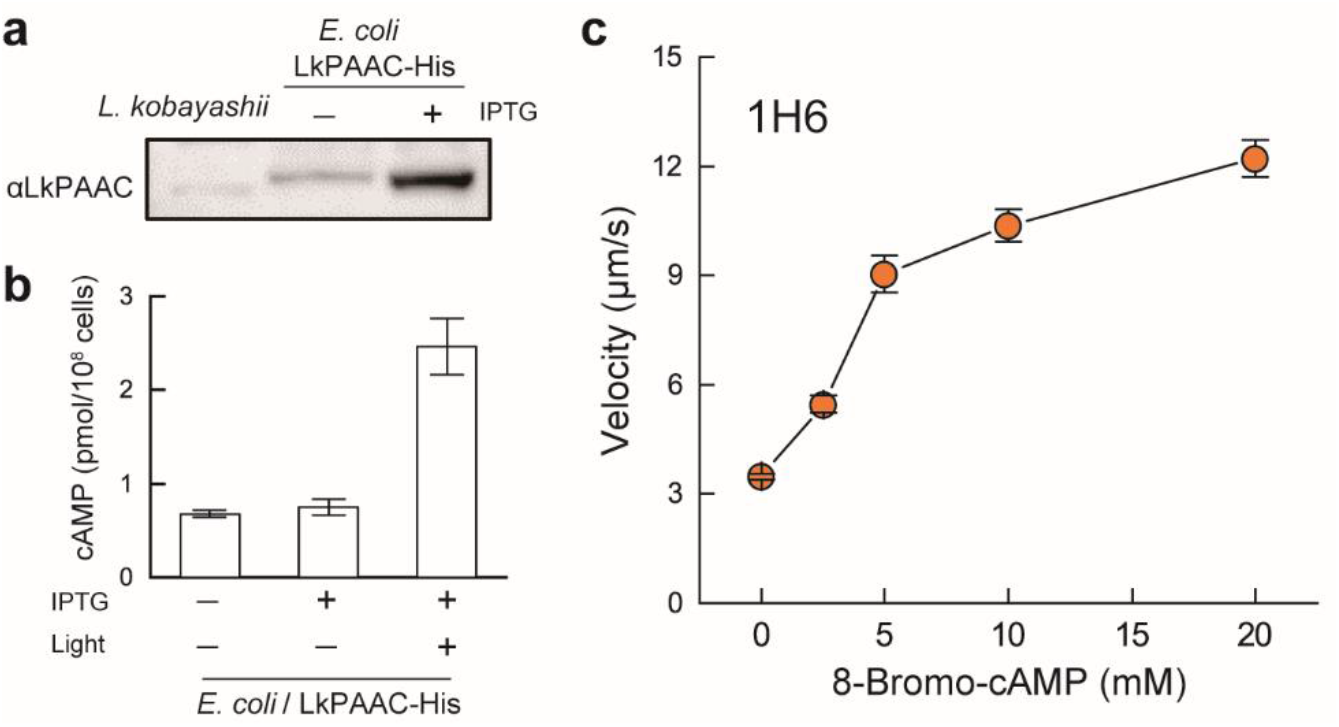
Activity and localization of LkPAAC. (a) Immunoblotting of whole-cell lysates of *L. kobayashii* (1.5× 10^8^ cells) and *E. coli* expressing LkPAAC-His (2.0× 10^7^ cells) with anti-LkPAAC antiserum. Since long-time exposure was required for detecting the band of *L. kobayashii*, the *E. coli* sample was diluted at 1:10 for observing the band of the overexpressed proteins clearly. (b) Light-dependent synthesis of cAMP in *E. coli* carrying the codon-optimized *lkPAAC* gene. The average values and standard deviations were determined by three independent experiments. (c) Effect of externally supplemented membrane-permeable cAMP analog (8-Bromo-cAMP) on the 1H6 swimming. The average values and standard errors were determined by three independent experiments (ca. 100 cells were measured in total in each concentration).

### Bi-polar localization of the photoresponsive cyclase

We examined the localization of LkPAAC in the cell body by labeling the protein with GFP. The GFP labeling did not affect the photoresponsivity of the bacterium (Supplementary Fig. 7). Epi-fluorescent microscopy showed the localization of LkPAAC-GFP at both ends of the cell body (Fig. 4a). Based on the fluorescent intensity of single AcGFP molecules, we estimated that 5.5 ± 3.8 molecules of LkPAAC (n = 46 poles) are localized at one pole (Fig. 4b) while those in the membrane pool diffused along the cell body (Movie 3). Although the increase of cAMP concentration in *L. kobayashii* was likely to be below the detection limit (Supplementary Fig. 6), the bi-polar localization of LkPAAC could condense cAMP near the flagellar motor and realizes rapid response to light.

**Figure 4.**
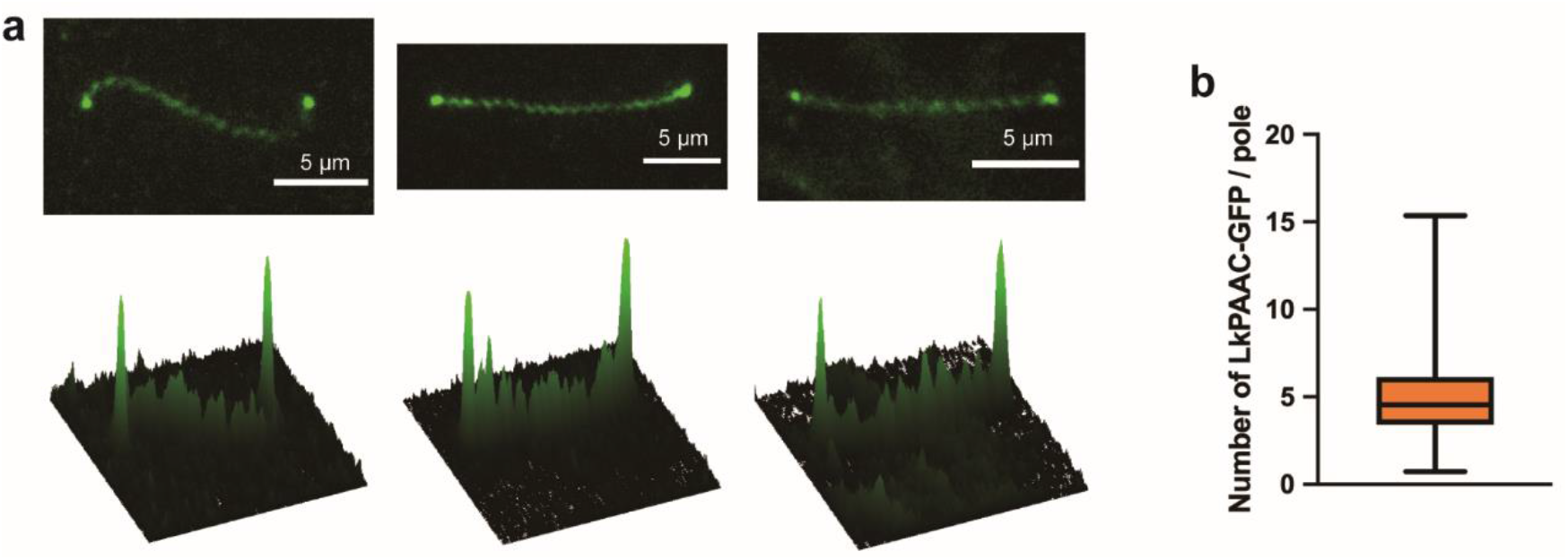
Subcellular localization of LkPAAC. (a) Fluorescent images of LkPAAC-GFP expressed in the 1H6 strain (upper) and 2D intensity profiles (lower). (b) Box plot of the number of LkPAAC-GFP molecules localized at one end of the cell body. The lower and upper box boundaries are 25th and 75th percentiles, respectively. The line in the middle of the box shows the median number. The lower and upper error lines indicate the smallest and largest values, respectively. 46 cell poles were analyzed.

## Discussion

Organisms react to light through various sensory systems, leading to vision, signaling, and energetic activities. For example, the retinal-binding membrane proteins rhodopsins play a crucial role in vision, ion pumping, and microbial taxis^25,26^; the flavin-binding proteins cryptochromes are involved in the growth of plants and the circadian clock of animals^27^. Investigating such light-driven proteins deepens our understanding of essential cellular activities. In addition, they have great potential for application as an optogenetic tool, enabling energetic or signaling modulation in live cells under arbitral spatial and temporal conditions. The adenylyl cyclase discovered in the soil spirochete modulated the swimming pattern through cAMP synthesized in a light-dependent manner. Although cAMP signaling has been known to be involved in pilus/flagella biogenesis and biofilm formation in bacteria, they are slow responses through gene regulation^10,12,28^. Our finding of subsecond-order stimulation of bacterial swimming by cAMP shows a novel role of the nucleotide second messenger. Subcellular localization of the LkPAAC near the flagellar motors implies a high local concentration of cAMP, offering a rapid and sensitive response of the flagellar rotation.

The genus *Leptospira* comprises pathogenic and saprophytic species^8^. The species that belong to the saprophytic clade S2 containing *L. kobayashii* can respond to light, whereas one of the major pathogenic species *L. interrogans* swims even in the dark and does not react to light despite carrying the homologous gene (Supplementary Fig. 8). Since the LkPAAC-deficient mutant retains motility but cannot migrate (Movie 2), cAMP-dependent swimming could be important for exploring environments of the free-living species. In contrast, the pathogens’ motility in irrelevant to light is significant for migrating within the host body^29^. Interestingly, a search of the amino-acid sequence database showed that many microbial species have a gene homologous with LkPAAC in their genome (Supplementary Fig. 9). Although LkPAAC would not be involved in photoresponsive motility in all bacteria, cAMP synthesized by LkPAAC may be used in some signal transductions. Adenylyl cyclases are categorized into six classes, e.g., class I has been found in gamma-proteobacteria, and class II is used by pathogenic bacteria, such as *Bacillus anthracis* and *P. aeruginosa*^30^. Since PACs that have functional similarity to LkPAAC belong to a universal class III used by many species of eukaryotes (e.g., fungi and protozoa) and prokaryotes (e.g., eubacteria and archaea), successful expression of functional LkPAAC in other bacterial species is the crucial first step for optogenetic application. Understanding the molecular basis of the fast photosensory of LkPAAC and cAMP signaling will be a future subject.

## Methods

### Bacteria and media

*Leptospira* spp. were grown in enriched Ellinghausen-McCullough-Johnson-Harris (EMJH) liquid medium (BD Difco, NJ, USA) at 30°C for 2 to 4 days (depends on the strains) until the log-phase. *E. coli* strains C41(DE3) and β2163 were grown in L-broth and LB medium supplemented with 0.3 mM diaminopimelate, respectively.

### Screening of a photoresponsivity-deficient mutant by random transposon mutagenesis

Random insertion mutagenesis of *L. kobayashii* strain E30 using *Himar1* transposon was conducted as described previously^31,32^. By conjugation with *E. coli* β2163 carrying pCjTKS2^32^, about 2400 transconjugant colonies were observed on plates of EMJH agar (1% agar) containing kanamycin at a final concentration of 25 μg/ml. Each transconjugant was independently inoculated to 150 μl of liquid EMJH containing kanamycin and grown at 30°C for 4 days. The grown bacteria were observed using a dark-field microscope (BX50, mercury lamp, 10x objective, dry condenser; Olympus) for screening photoresponsivity-deficient mutants. The transposon insertion site was identified using the semi-random PCR technique^32^.

For the complementation of photoresponsive-deficient mutant 1H6, LPTSP3_g09850 and/or *lkPAAC* (LPTSP3_g09840) genes were expressed under the *flgB* promoter^33,34^. The *flgB* promoter region was amplified as previously described^29^, and the LPTSP3_g09850/*lkPAAC* genes, *lkPAAC* gene, LPTSP3_g09850 gene with FLAG tag, and *lkPAAC* with FLAG tag were amplified from genomic DNA of the *L. kobayashii* E30, and the amplified products were cloned into the *Sal*I-digested pCjSpLe94^35^ by NEBuilder HiFi DNA Assembly cloning (New England BioLabs). For the LkPAAC with GFP joined by a flexible (GGGGS)3 linker, the *lkPAAC* gene was amplified as described above, and *gfp* was amplified from pAcGFP1 (Clontech) using the primer containing the (GGGGS)3 linker sequence, and the amplified products were cloned into the *Sal*I-digested pCjSpLe94 as described above. Primers used in this study are listed in Supplementary Table 1.

### Immunoblotting experiments

About 1.5 × 10^8^ leptospiral cells suspended in SDS-PAGE sample buffer were subjected to 5%-20% SDS-PAGE and Western blotting. The blot was incubated with antisera raised against the peptide fragment of LkPAAC (NH2-LSWADRTDSIYIWK-COOH) and FlaA2^31^ or monoclonal antibody for FLAG tag.

### Motility assay

*Leptospira* culture was diluted 1:20 into a fresh EMJH and was infused into a flow chamber composed of a coverslip (upper side) and a glass slide (bottom side). To examine the effect of membrane-permeable cAMP and cGMP on swimming, 8-Bromo-cAMP and 8-Bromo-cGMP dissolved in 10 mM Tris-HCl (pH7.0) were added to *L. kobayashii* cells suspended with the same buffer. Motility of leptospires was observed under a dark-field microscope (BX53, Splan 40×, NA 0.75; Olympus, Tokyo, Japan) equipped with a halogen lamp and were recorded by a supersensitive charge-coupled device (CCD) camera (WAT-910HX, Watec Co., Yamagata, Japan) at a frame rate of 30 frames per second. Swimming trajectories and velocities of individual cells were analyzed using ImageJ software (National Institutes of Health, MD, USA) and VBA-based macros programmed in Microsoft Excel (Microsoft, WA, USA)^14^. Light intensity and wavelength were adjusted with neutral density (ND) filters and bandpass filters (FF01-488/50-25 for blue, FF01-550/49-25 for green, FF01-650/54-25 for red; Semrock), respectively. Light intensity was measured using an illuminometer (CHE-LT1, Sanwa Supply INC.)

### Observation of LkPAAC-AcGFP1 subcellular localization

Fluorescence of LkPAAC-AcGFP was observed using an inverted fluorescence microscope (IX-83, Olympus) with a 100× oil immersion objective lens (UPLSAPO100XO, NA 1.4, Olympus) and an sCMOS camera (Prime95B, Photometrics). AcGFP was excited by a 130 W mercury light source system (U-HGLGPS, Olympus) with a fluorescence mirror unit U-FGFP (Excitation BP 460–480; Emission BP 495–540, Olympus). Fluorescence image processing was performed with the ImageJ version 1.53 software (National Institutes of Health).

The number of LkPAAC-AcGFP1 at the cell pole was estimated by comparing the fluorescence spot intensity at the pole with the fluorescence intensity of a single His-AcGFP1 molecule as previously reported^36^. His-AcGFP1 was purified from *E. coli* C41(DE3) cells carrying pET19b/ His-AcGFP1 using Ni-NTA agarose (Fujifilm Wako). 10 pg/ml of His-AcGFP1 solution was applied to a coverslip washed by 0.1 M KOH and observed by fluorescence microscopy. In the fluorescent images, a rectangular mask for the fluorescent spot of 30×30 pixels was applied to the ROI (region of interest). We defined the spot intensity as the sum of all pixel values within the rectangular mask after subtracting the total background intensity from each pixel value. The number of LkPAAC-AcGFP1 per pole was estimated as the intensity of the cell pole divided by the average intensity of a single His-AcGFP1 molecule.

### cAMP ELISA assay

A cAMP ELISA kit (ADI-900-066, Enzo Life Sciences) was used to determine intracellular cAMP levels following the manufacturer’s instructions. *E. coli* C41(DE3) cells carrying pET22b/ LkPAAC-His were grown in L-broth containing 100 μg/mL ampicillin with or without 1 mM IPTG at 30°C for 5 hours with shaking. The codon usages of the LkPAAC sequence were optimized to those of *E. coli* for efficient protein expression. *L. kobayashii* cells were grown at 30°C for 4 days in EMJH liquid medium until the late-exponential phase. Cells were photo-stimulated with white LED for 3 min and subsequently lysed. Cell lysates were used to calculate cAMP values. The intracellular cAMP concentrations of *E. coli* and *Leptospira* cells were normalized to OD600 and OD420, respectively.

## Acknowledgments

We thank Dr. E. Isogai (Tohoku University) and Dr. H. Yoneyama (Tohoku University) for the experiment reagents and the insightful discussion. This work was supported by the JSPS KAKENHI Grant number 18J10834 to J.X., 19K07571 to N.K., 15H01307 to S.N., and 20H05524 to S.N. This work was also supported by JST PRESTO Grant Number JPMJPR204B to Y.V.M.

## Author contributions

J.X., N.K., Y.V.M., R.O., T.M., and S.N. planned the project; J.X., N.K. Y.V.M. R.O., T.M., and S.N. carried out the experiments; Y.V.N. and S.N. set up the optical system and programs for data analysis; J.X., Y.V.M., N.K., R.O., and S.N. analyzed the data; J.X., N.K., Y.V.N., and S.N. wrote the paper.

## Competing interests

The authors declare that they have no competing interests.

## Data availability

The data supporting the findings of this study are available from the corresponding author upon request.

## Notes

### Competing Interest Statement

The authors have declared no competing interest.

