## Supplementary Information for "Light-dependent synthesis of a nucleotide second messenger controls bacterial motility"

Supplementary Table 1. Primer sequences used in this study.

Supplementary Fig. 1. Morphology of *Leptospira*.

Supplementary Fig. 2. Quantification of the light-dependent change in motility pattern using mean square displacement (MSD) analysis.

Supplementary Fig. 3. Light-intensity dependence of the *L. kobayashii* responsivity.

Supplementary Fig. 4. Effect of the LPTSP3_g09840 gene complementation on the photoresponsivity of the 1H6 mutant

Supplementary Fig. 5. Sequence homology between BDA78054 (LkPAAC) and photoactivated adenylyl cyclases

Supplementary Fig. 6. Enzyme activity of BDA78054 (LkPAAC)

Supplementary Fig. 7. Effect of GFP labeling on photoresponsivity of *L. kobayashii*.

Supplementary Fig. 8. Photoresponsivity of the relative species

Supplementary Fig. 9. Conservation of LkPAAC in prokaryotes.

Movie 1. Photoresponsivity of *L. kobayashii* at 400-750 nm. The cells were observed in dark first (Light OFF), and then exposed to light (Light ON).

Movie 2. Photoresponsivity of 1H6 mutant. The cells were observed at the “Bright” condition (see Supplementary Fig. 2)

Movie 3. 1H6 mutant expressing GFP-labeled LkPAAC. The cell poles are indicated by white arrows. The movie was recorded at 2 frames per second and replayed at 10 times speed.

**Supplementary Table 1.** Primer sequences used in this study

| Target | Primer sequence (5´→ 3´) | |
|---|---|---|
| | Forward | Reverse |
| LPTSP3_g09850 | TAAATGAGGGAGGTTTCCATATGAAAAGATTACATCCGTCAGG | GCGAGGCTGGCCGGCGTCGATCATAATGTTTGAATGGAAACGAAAGCTC |
| *lkPAAC* (LPTSP3_g09840) | TAAATGAGGGAGGTTTCCATATGATAGACTTAAATTACATACTCGC | GCGAGGCTGGCCGGCGTCGATTAAGGTTTCCAAATATAAATGGAATCCG |
| LPTSP3_g09850/*lkPAAC* | TAAATGAGGGAGGTTTCCATATGAAAAGATTACATCCGTCAGG | GCGAGGCTGGCCGGCGTCGATTAAGGTTTCCAAATATAAATGGAATCCG |
| LPTSP3_g09850 with FLAG tag | TAAATGAGGGAGGTTTCCATATGAAAAGATTACATCCGTCAGG | GCGAGGCTGGCCGGCGTCGATCACTTGTCATCGTCATCCTTGTAGT<br>CTAATGTTTGAATGGAAACGA |
| *lkPAAC* with FLAG tag | TAAATGAGGGAGGTTTCCATATGATAGACTTAAATTACATACTCGC | GCGAGGCTGGCCGGCGTCGATCACTTGTCATCGTCATCCTTGTAGT<br>CAGGTTTCCAAATATAAATGG |
| *lkPAAC* for AcGFP fusion | TAAATGAGGGAGGTTTCCATATGATAGACTTAAATTACATACTCGC | CCGCCGGAACCGCCTCCACCAGGTTTCCAAATATAAATGGAATCCG |
| AcGFP+(GGGGS)$_3$ linker | GGTGGAGGCGGTTCCGGCGGAGGTGGCTCCGGCGGTGGCG<br>GATCCATGACCATGATTACGCCAAGC | GCTGGCCGGCGTCGATCACTTGTACAGCTCATCCATG |

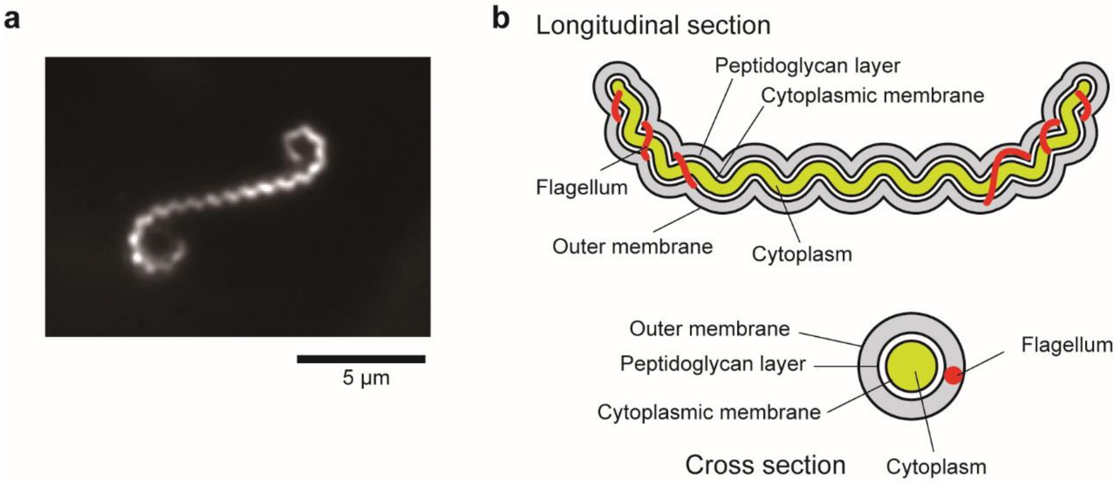


**Supplementary Fig. 1.** (a) *Leptospira kobayashii* observed by dark-field microscopy. (b) Schematic diagram depicting the structure of *Leptospira* spp.

**Quantification of the light-dependent change in motility pattern using mean square displacement (MSD) analysis.**

Consider two particles moving to the right (positive) or left (negative) with time as shown in Supplementary Fig. 2a. The upper particle (gray) moves to both positive and negative directions with the same possibility (*P*), which is so-called free diffusion. In contrast, the movement of the lower one (red) is biased to the positive side. The bead displacements were determined by comparing the assumed probabilities and automatically generated random numbers (rnd); e.g., If P+ > rnd, the bead moves in a positive direction. The result of the simulation is shown in the right panel. The one-dimensional MSD of such particles during Δt is calculated by $MSD(\Delta t) = \langle (x_{i+\Delta t} - x_i)^2 \rangle$, where $x_i$ is the particle position at time $i$. The time vs MSD plot of the diffusive motion shows linear relation to time, whereas that of the biased movement shows a quadratic curve (Supplementary Fig. 2b, left). Therefore, double-logarithmic plots show linear lines with h slopes of ~1 and ~2 (Supplementary Fig. 2b, right).

We used MSD analysis to quantify the light-dependent motility of *L. kobayashii*. Supplementary Fig. 2c are kymographs of the bacteria moving in dark and bright conditions. The MSD values were computed using data of temporal cell positions, and example traces obtained from about 50 cells are shown in Supplementary Fig. 2d. The double-logarithmic traces and the slopes determined by line fitting to individual traces of all measured samples revealed that most of the bacteria moving in bright have a slope >1, whereas those in the dark have <1 (Supplementary Fig. 2e). These results indicate that the bacterial migration is strongly suppressed while the cell is rotating in the dark, and the movement is instantly directed upon light exposure.

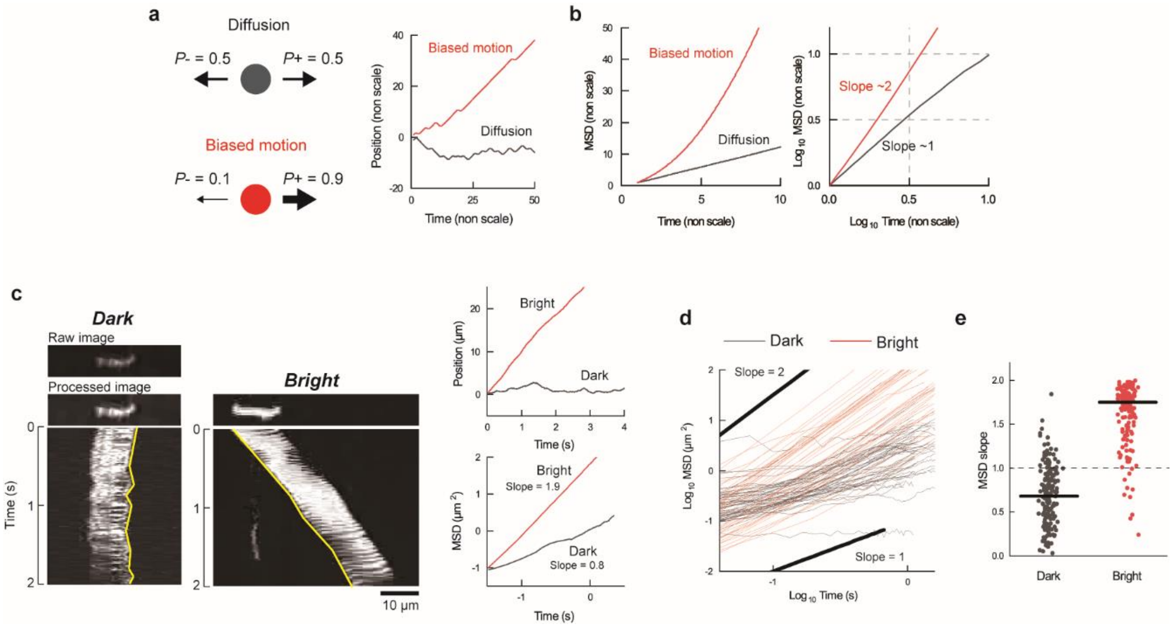


**Supplementary Fig. 2.** (a) Simulation of diffusive (gray) and biased (red) motions. (b) Time plots of MSD obtained from the simulated bead displacement in ***a***. (c) Kymographs of *L. kobayashii* cells moving in dark and bright conditions. The brightness of video images recorded in the dark was adjusted so that the cell position could be determined by ImageJ software. The yellow lines indicate the cell movement. The right panels are time courses of cell positions (upper) and MSD (lower) of the bacteria shown on the left. (d) Example MSD time plots; 59 and 45 traces (cells) are shown for dark and bright, respectively. (e) Summary of the values of MSD slope obtained by line fitting to individual paths; 189 and 150 cells were analyzed in dark and bright, respectively. The horizontal bars indicate the median: 0.68 for dark and 1.75 for bright.

### Light-intensity dependence of the *L. kobayashii* responsivity.

We tested the dependence of the light-responsive bacterium *L. kobayashii* on the light intensity. The light intensity was measured using an illuminometer (CHE-LT1, Sanwa Supply INC.). Supplementary Fig. 3a-c show that the swimming velocity and MSD slope (see Supplementary Fig. 2) of the bacterial population increase with light intensity. Although the velocity is decreased at 14640 lux, the result that the MSD slope is still close to 2 implies uncertain photodamage to the bacteria.

As discussed in Supplementary Fig. 2, time courses of the individual bacterial movements show that light stimulation affects the frequency of back-and-forth motion (Supplementary Fig. 3d). The sequential images recorded at 105 lux show that the bacterium alternates swimming (the periods shown in red in Supplementary Fig.3e) and rotation (those shown in gray), and the rotation period could be interpreted as “tumbling” observed in peritrichous bacteria such as *Escherichia coli*. Exponential distributions were obtained by measuring the time for unidirectional swimming (the left panels of Supplementary Fig.3f), and the rate constants of the transition from swimming to tumbling ($k_{Sw\rightarrow Rot}$) were determined by exponential fitting to the experimental histograms using $exp(-k_{Sw\rightarrow Rot}t)$ (red lines are the results of fitting). The reversal rate is decreased with increased light intensity (the right panel), indicating that the light stimulation suppresses the reversal event of the flagellar rotation. Swimming time determined as a reciprocal of $k_{Sw\rightarrow Rot}$ is extended with the light intensity (Supplementary Fig 3g), where the swimming time at 11 lux was plotted as zero because of no detectable swimming during observation.

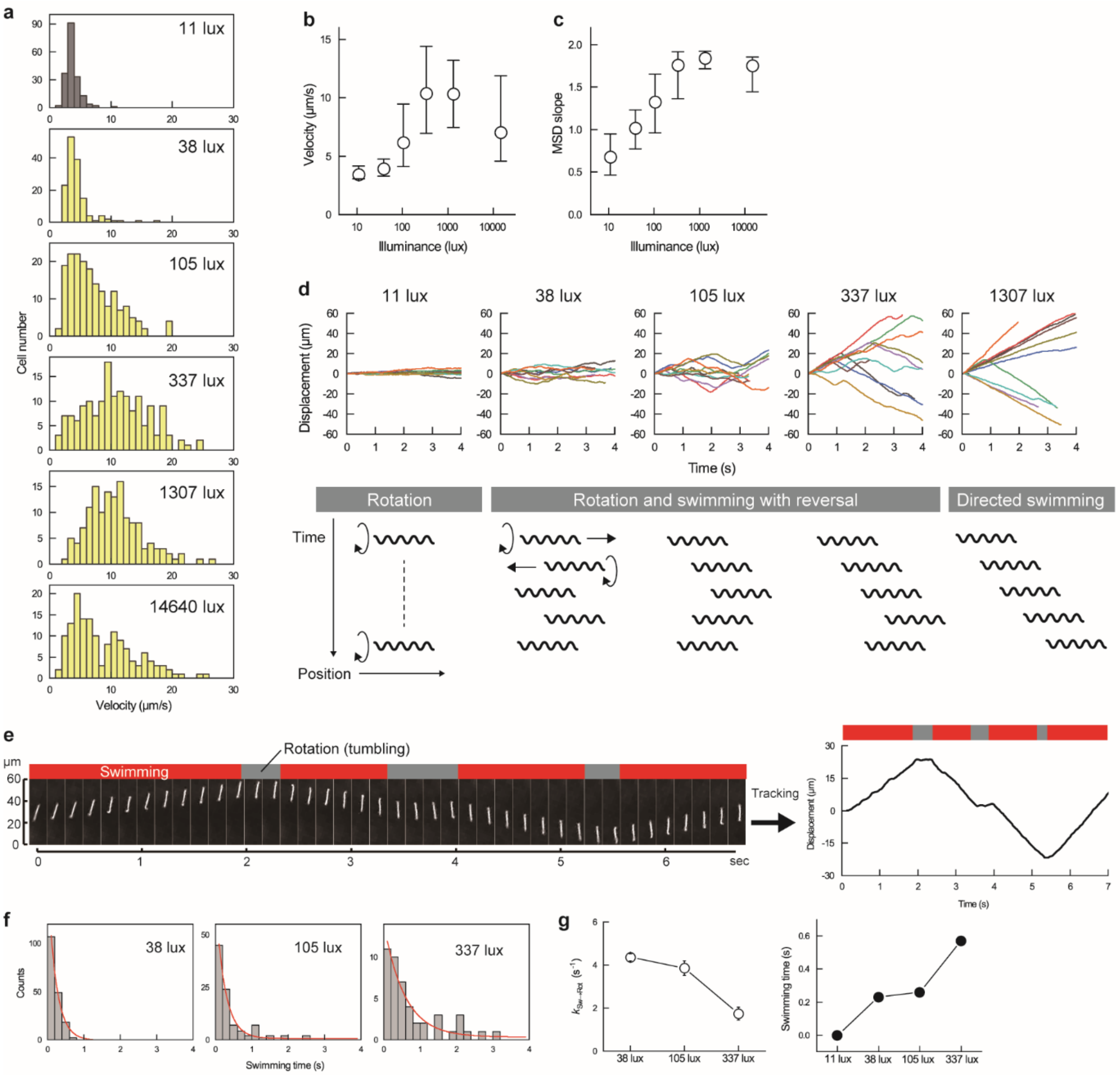


**Supplementary Fig. 3.** (a) Velocity distributions of *L. kobayashii* swimming in various light-intensity conditions. (b) and (c) show the 25th (lower whiskers), 50th (open circles), and 75th (upper whiskers) percentiles of velocity and MSD slope (see Supplementary Fig 2), respectively. Illuminances indicated in the histograms were average values of five measurements. Each histogram was obtained by three independent experiments, and 183 cells for 11 lux, 145 cells for 38 lux, 179 cells for 105 lux, 165 cells for 337 lux, 141 cells for 1307 lx, and 150 cells for 14640 lx were measured in total. The data of 11 lux and 1307 lux were the same as "Dark" and "Bright" in Fig 1. (d) The upper panels are time courses of bacterial movements. Trajectories obtained from 10 bacteria were shown in each condition. The lower cartoons schematically explain the bacterial movements observed in each light intensity. (e) Sequential images of a *L. kobayashii* cell swimming at the light intensity of 105 lux (left), and the results of cell tracking (right). (f) Distributions of the swimming time. The red curves are the results of the exponential fitting. (g) The rate constants of the transition from swimming to rotation determined by the fitting in ***f*** (left; error bars are the standard error of fitting) and the reciprocals of the rate constants, i.e., the swimming time (right).

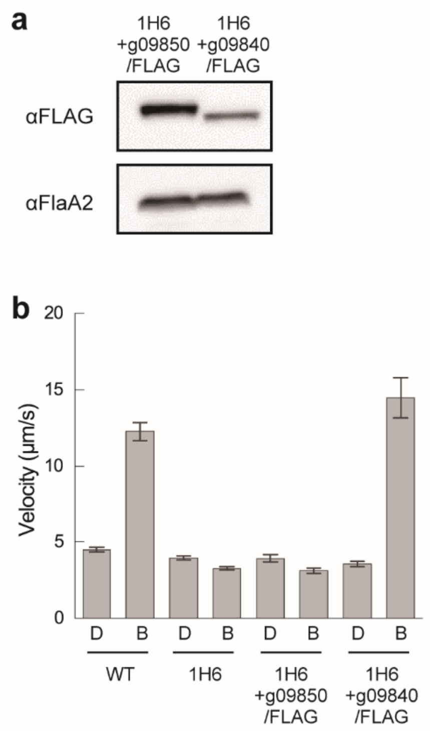


**Supplementary Fig. 4.** Effect of the LPTSP3_g09840 gene complementation on the photoresponsivity of the 1H6 mutant. (a) Immunoblotting of whole-cell lysates from the 1H6 strain complemented by LPTSP3_g09850 gene with FLAG-tag or LPTSP3_g09840 gene with FLAG-tag. The anti-FlaA2 antiserum and monoclonal antibody for FLAG tag were used. (b) Swimming velocities measured in Dark (D) and Bright (B) conditions. Average values and standard errors are shown (n= 75 cells for WT dark, 193 cells for WT bright, 85 cells for 1H6 dark, 169 cells for 1H6 bright, 38 cells for g09850/FLAG dark, 70 cells for g09850/FLAG bright, 45 cells for g09840/FLAG dark, and 39 cells for g09840/FLAG bright).

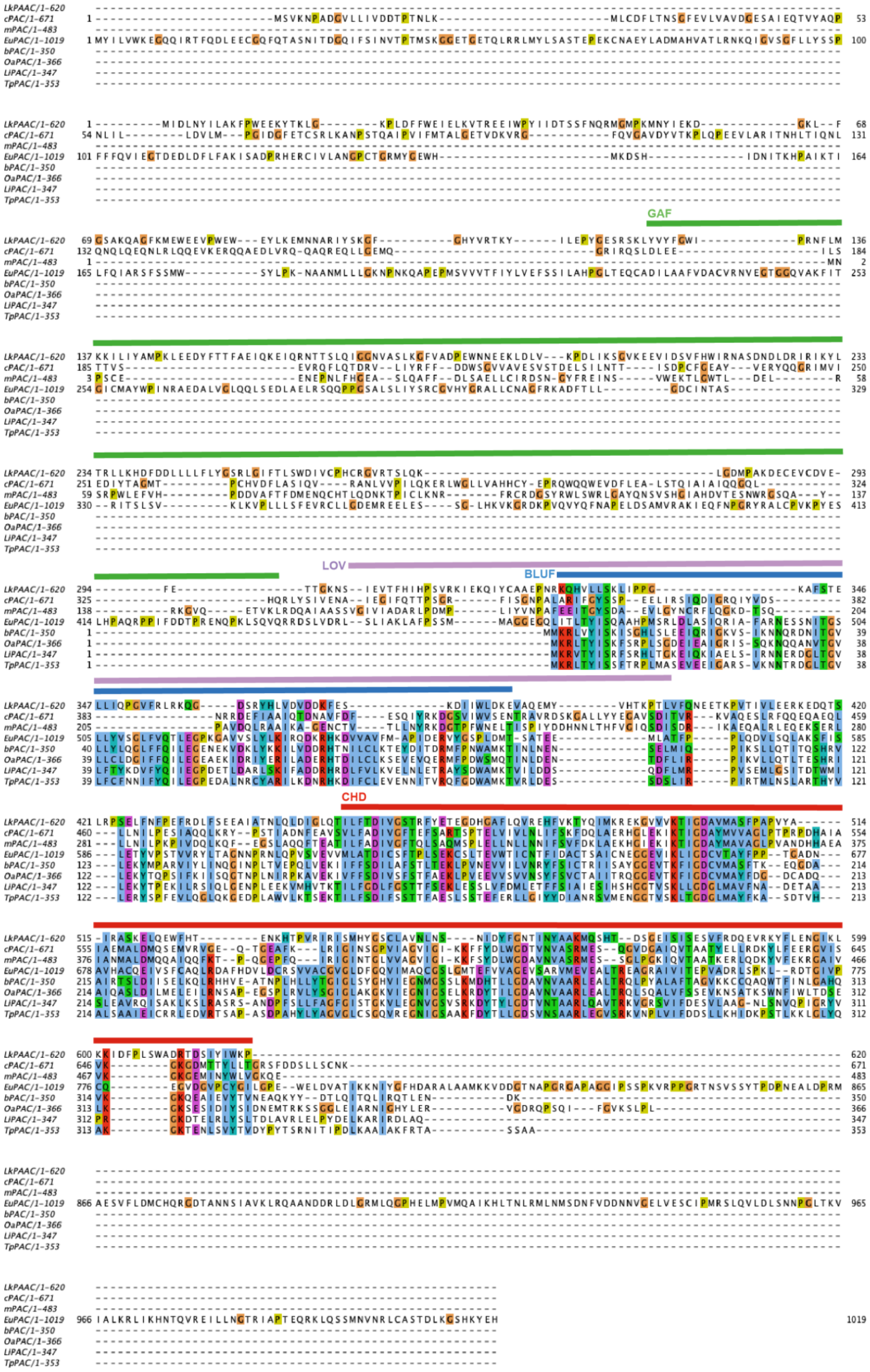
GAF
LOV
BLUF
CHD
LkPAAC/1-620
cPAC/1-671
mPAC/1-483
EuPAC/1-1019
bPAC/1-350
OaPAC/1-366
LiPAC/1-347
TpPAC/1-353

**Supplementary Fig. 5.** Alignment of amino acid sequences of LkPAAC with known functional PACs. The source organisms are *Microcoleus* sp. *PCC 7113* (cPAC)[1], *Microcoleus chthonoplastes* PCC7420 (mPAC)[2], *Euglena gracis* (EuPAC)[3], *Negleria grubei* (NgPAC2)[4], *Baggiatoa* spp. (bPAC)[5], *Oscillatoria acuminata* (OaPAC)[6], *Leptonema illini* (LiPAC)[7], and *Turneriella parva* (TpPAC)[8]. Multiple sequence alignment was analyzed and visualized using Jalview[9]. Only cPAC contains GAF domain (green bar); only mPAC contains LOV domain (purple bar); EuPAC, bPAC, OaPAC, LiPAC, and TpPAC have BLUF domain (blue bar) as a sensor. LkPAAC contains the cyclase homology domain (CHD, indicated by the red bar) but has neither BLUF nor LOV domain.

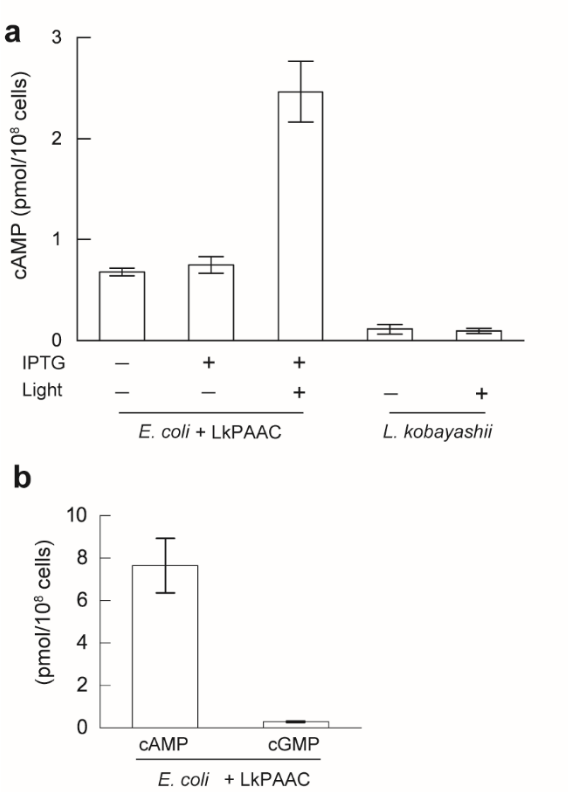


**Supplementary Fig. 6.** (a) The cAMP concentration measured in the *E. coli* carrying the codon-optimized LkPAAC (*E. coli* + LkPAAC) and *L. kobayashii* with or without light exposure for 3 min. (b) The cAMP and cGMP concentration measured in *E. coli* + LkPAAC after light exposure for 30 min. Average values and standard deviations of three independent experiments are shown.

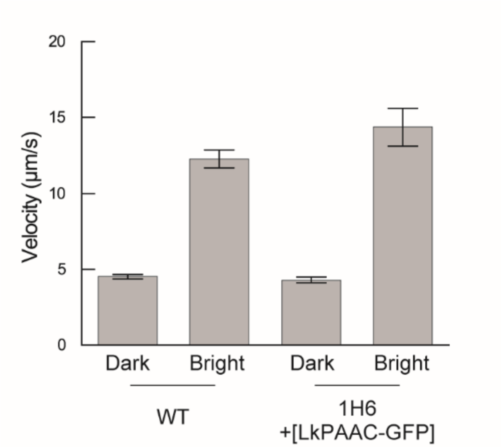


**Supplementary Fig. 7.** Effect of GFP labeling on photoresponsivity of *L. kobayashii.* The data of WT are the same as those shown in Fig 1d (green). For 1H6, 46 cells and 55 cells were measured in dark and bright with the green filter, respectively. Average values and standard errors are shown.

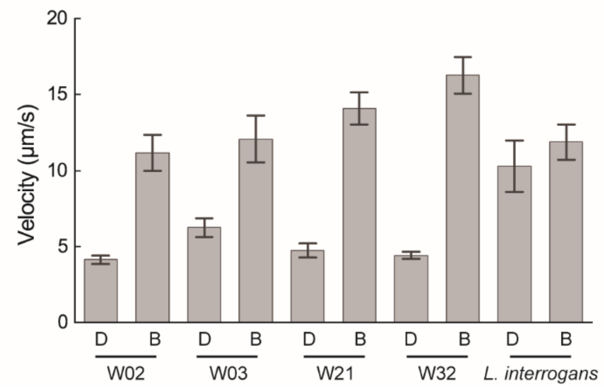


**Supplementary Fig. 8.** Photoresponsivity of the *Leptospira* species that are phylogenetically related to *L. kobayashii*. The strains were isolated from water and classified into the saprophytic clade including *L. kobayashii* based on the 16S ribosomal RNA sequence[10]. *L. interrogans* is one of the most important pathogenic species, causing the worldwide zoonosis leptospirosis. The swimming velocity was measured in dark (D) and bright (B), and average values and standard errors determined from data of 20–50 cells are shown.

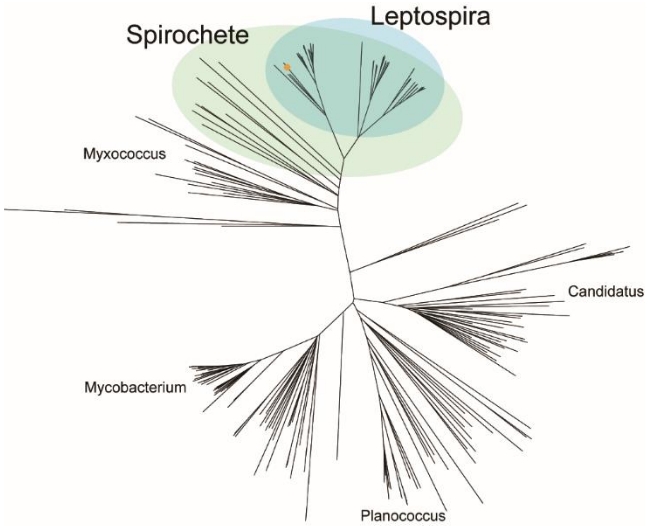


**Supplementary Fig. 9.** Conservation of LkPAAC in prokaryotes. The orange dot indicates *L. kobayashii*. Homologous genes of LkPAAC were searched by PSI-BLAST program (NCBI). Maximum likelihood phylogenetic trees were constructed and visualized using MEGA X[11] and iTOL program[12].
